# Fingerprint localisation for fine-scale wildlife tracking using automated radio telemetry

**DOI:** 10.1101/2024.02.15.580447

**Authors:** Chris Tyson, Rita Fragueira, Emilio Sansano-Sansano, Hui Yu, Marc Naguib

**Affiliations:** Behavioural Ecology Group, Wageningen University & Research, Wageningen, The Netherlands; Institute of New Imaging Technologies, Universitat Jaume I, Castellón, Spain; Experimental Zoology Group, Wageningen University & Research, Wageningen, The Netherlands

**Keywords:** automated radio telemetry, fingerprint localisation, multilateration, movement ecology

## Abstract

1: Automated radio telemetry systems are among the most widely applicable method for tracking wildlife at fine spatiotemporal scales. These systems are increasingly applying large networks of simple receivers to detect radio signal strength (RSS) from radio tagged individuals. Analytical methods to derive position estimates using RSS localisation, however, are relatively underdeveloped in the field of wildlife tracking. Here, we apply approaches from indoor positioning systems to develop a new method, radio fingerprinting, for localizing radio tagged animals in structurally complex, outdoor environments. This method characterizes the RSS patterns at known locations to generate a radio map of the study area, which is then used to predict the location of new RSS patterns.

2: We conducted field tests to evaluate the localisation accuracy of radio fingerprinting relative to multilateration, a commonly used method for RSS localisation. To do so, we established an experimental receiver network covering multiple habitat types and compared the localisation accuracy of radio fingerprinting to multilateration. Additionally, we evaluated how a variety of features characteristic of typical tracking datasets affected the accuracy of both methods.

3: While both methods had a similar median error (∼30 m), the fingerprint localisation method offered several advantages over multilateration. Multilateration localisation estimates were highly affected by missed detections from the nearest receiver to the test point, which occurred in 30% of cases. In these cases, the median error was 97 m, over a 3-fold increase. Distance to the nearest receiver also biased multilateration estimates with error increasing as the distance increased. Additionally, errors from multilateration estimates were higher in more densely vegetated areas. In contrast, fingerprinting position estimates were largely robust to each of these scenarios.

4: Automated radio telemetry enables the fine-scale, continuous tracking of range-resident animals. We present radio fingerprinting as a localisation method in outdoor environments where standard localisation methods may be inadequate. This approach can be applied to any automated radio telemetry hardware and to any study system where a radio map can be generated.

## Introduction

Advances in tracking technologies have given wildlife researchers unprecedented abilities to study the movement of animals in the wild. As movement is an integral component of many biological processes, tracking an animal’s location through time can yield insights into diverse topics such as life history, space use, communication, and social behaviour (Kays et al., 2015; Nathan et al., 2008). Fundamental trade-offs between tracking tag size and sampling duration and rate, due to energy requirements, however, requires that researchers must choose a tracking method that is suitable for the species and the question of interest (Nathan et al., 2022; Snijders et al., 2017). For example, while satellite tracking (e.g., GPS tags) can offer high spatial accuracy, these tags are energy demanding and so cannot operate at a high frequency over extended periods without becoming prohibitively heavy for use on small animals (Nathan et al., 2022). In contrast, lightweight, solar-powered digital radio tags are suitable for most size birds and mammals, and the relative affordability of these tags allows multiple individuals to be simultaneously tracked (Bircher et al., 2020; Scardamaglia et al., 2022; Shizuka et al., 2022). Historically, however, radio telemetry required tagged individuals to be manually followed individually with hand-held receivers, which limited concurrent tracking of groups of animals as well as the spatial extent and continuity of these radio tracking studies (Kenward, 2001).

Automated radio tracking systems (ARTS) have provided a major leap forward for radio telemetry studies by enabling multiple radio tagged individuals to be simultaneously and continuously monitored over areas bounded only by the presence of radio receivers (Mennill et al., 2012). Various ARTS designs exist, but the general framework is a network of receivers that detect radio signals from tags and record the relevant information (e.g., signal strength, time of arrival, or signal bearing).

Simultaneous detections by multiple (minimally three) receivers can then be analysed to estimate the location of the tag (Beardsworth et al., 2022; Kays et al., 2011). A drawback of some of these systems is that they can be technically challenging to operate. For example, time of arrival systems require highly accurate clocks (Maccurdy et al., 2009) and function poorly in areas without line-of-sight or with extensive elevation gradients. As an alternative, the use of simple receivers with omni-directional antennas to measure received signal strength (RSS) is being rapidly adopted as an affordable and accessible ARTS for monitoring small, range-resident animals (Krull et al., 2018; Paxton et al., 2022; Wallace et al., 2022). Yet, while the tracking technology has developed in recent years the analytical methods to make optimal use of RSS-based localisation in wildlife studies has lagged behind the technological advances (but see Paxton et al., 2022).

Indoor positioning systems, which are used where satellite-based localisation is ineffective or impossible, commonly employ RSS-based localisation (Mendoza-Silva et al., 2019). One standard method for RSS-based localisation is multilateration. This method uses the known locations of time-synchronized receivers, each of which measures the RSS of a signalling tag. The RSS value registered by each receiver can then be modelled as a function of the logarithmic distance between the receiver and the signal, as well as by environmental properties (such as humidity) and hardware characteristics that also influence signal strength (Lee & Buehrer, 2011). Once the relationship between RSS and distance is determined, RSS values from three or more receivers can be used to estimate the location of a tag relative to the location of the receivers using multilateration (the general case of trilateration where three or more receivers are used). This method, as well as angulation methods, such as triangulation, are types of range-based localisation as they require knowing the positions of the receivers and the distance to the signal is explicitly modelled (Mendoza-Silva et al., 2019). Consequently, the accuracy of multilateration localisation depends critically on the relationship between RSS and distance, and uncertainty in this relationship will increase localisation error. As such, heterogenous, outdoor environments are especially challenging settings in which to use range-based localisation due to the complex sources of noise that are typically present, particularly in the form of structural features in the habitat, though additional data processing filters and calibrations steps can help to minimize the localisation error using multilateration (Paxton et al., 2022).

A widely used alternative to multilateration employed for RSS-based localisation for indoor positioning systems is RSS fingerprinting (He & Chan, 2016; Liu et al., 2007). The general approach of RSS fingerprinting is to characterize radio signal patterns at known locations in order to characterize the spatial variation of these patterns (Mendoza-Silva et al., 2019). As such, this method exploits features of the environment to characterize spatial patterns in RSS values and does not explicitly consider the distance between the signal and the receiver. For this reason, RSS fingerprinting is considered a range-free localisation method, in contrast to multilateration. RSS fingerprinting occurs in two stages. In the first stage, commonly called the offline stage, the signal pattern detected by each receiver (the fingerprint) is recorded at known positions (i.e., geographic coordinates) throughout the target tracking area. In this method, it is not necessary to know the locations of the receivers, though they are expected to remain constant. Taken together, these radio fingerprints comprise the radio map, which characterizes how a tag’s signal is detected by the receivers throughout the entire study area. For the offline stage, many methods exist to construct the radio map, spanning from relatively labour-intensive manual point-by-point calibration to more computationally demanding machine learning approaches in which the fingerprints are modelled (Jung, Moon, et al., 2017). In the second stage, the online stage, a new signal pattern is matched to similar fingerprints from the radio map to estimate the position of the signal. For the online stage, there are also many methods to estimate the position of a signal relative to the stored fingerprints, but in general there are three widely used classification approaches: deterministic, probabilistic, or machine learning (Jung, Moon, et al., 2017; Mendoza-Silva et al., 2019). While RSS fingerprinting is commonly applied for indoor positioning applications where environmental features are relatively static, it remains to be broadly evaluated as a localisation method for radio tracking in heterogenous, dynamic outdoor environments as used in tracking studies on the behaviour, ecology, and evolution of animals.

In this study, we evaluate the performance of RSS fingerprinting as a novel localisation method for fine-scale wildlife tracking using automated radio telemetry systems. To test this method, we used a large network of 49 receivers spanning a mosaic of habitat types. We then conducted manual point-by-point calibration throughout a test area to generate a radio map, which was used to estimate the tag locations using radio fingerprinting. Multilateration was also used to estimate test tag location and the localisation error of the two methods were compared under various scenarios characteristic of real-world radio tracking studies. As such, our overall objective in this study is to assess the effectiveness of RSS fingerprinting for wildlife tracking and to contrast this method against the more commonly used method of multilateration. In doing so, we aim to highlight a new localisation method for wildlife tracking studies in the rapidly growing field of automated radio telemetry and to provide an analytical workflow for utilizing this approach.

## Materials and Methods

### i. Field data collection

Testing was conducted at the Westerheide Wildlife Sanctuary (52.0162° N, 5.8412° E), the Netherlands, with permission of the local land managers, Geldersch Landschap & Kasteleen. The habitat in this area is a mosaic of managed beech forest, heather, livestock grazing fields, and new-growth pine. Within a 0.5 km^2^ region, we deployed 49 receivers in a 100 m triangular grid (Figure 1). For the receivers, we used the CTT Node v.2.0 (Cellular Tracking Technologies, Rio Grande, New Jersey, USA), which have an omni-directional, whip antenna (10 cm) that detects signals from 433 MHz radio tags and records the tag ID, detection time, and the RSS value. Detection data is stored on an internal micro-SD card and transmitted to a central base station along with the receiver ID. The receiver time is maintained by a real-time clock, which is routinely updated via satellite to maintain synchrony across the network. Receivers were placed approximately 2 m above ground and mounted to either PVC poles or small trees, depending on the location in the habitat.

**Figure 1.**
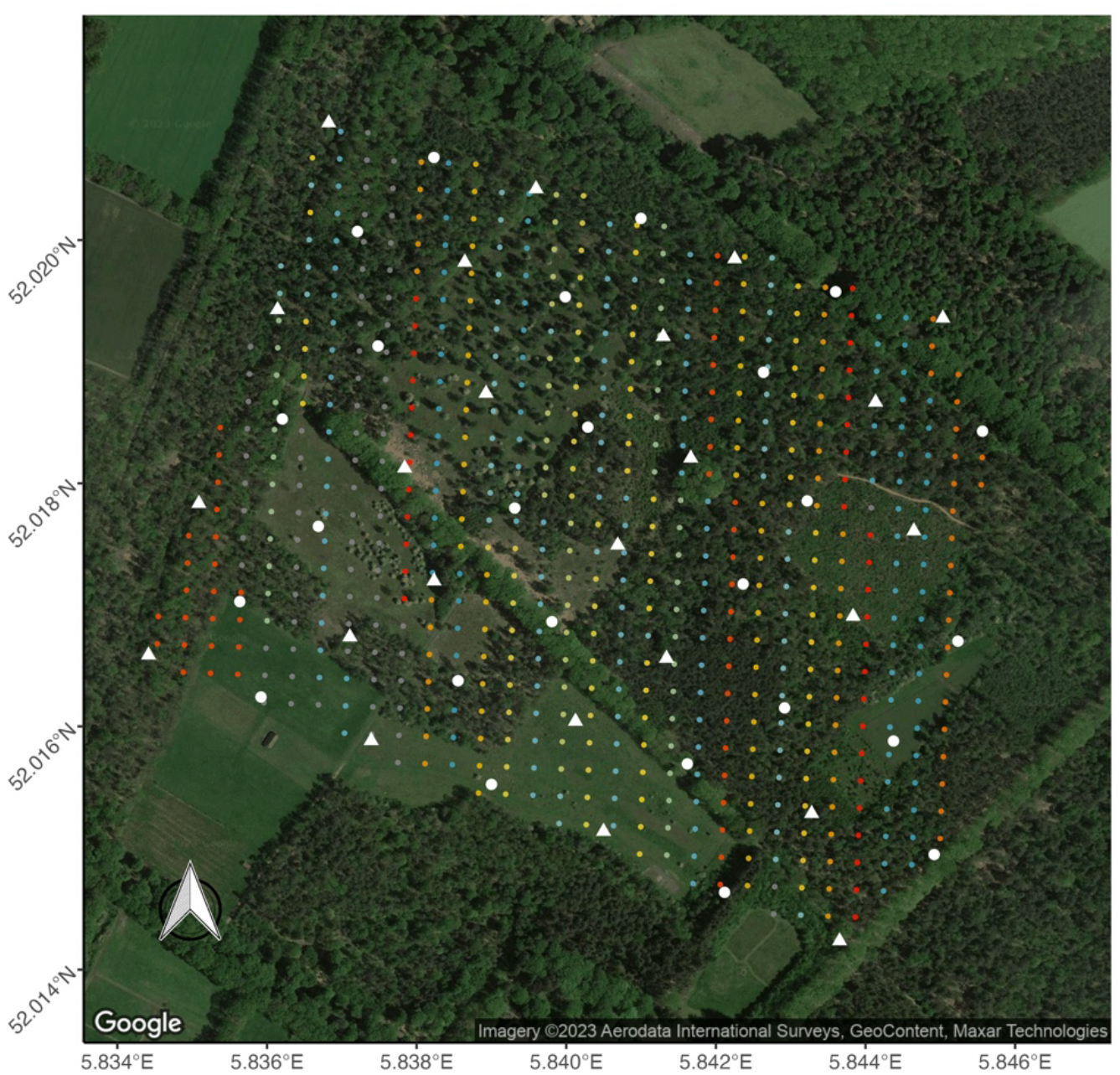
Map of the test area in Westerheide Wildlife Sanctuary, the Netherlands, showing the 49 receivers (white points) and varying habitat types within the receiver area. Receivers were spaced every 100 m on a triangular grid. Coloured circles show the test points (n = 532) spaced every 25 m on a square grid. Points are coloured according to the set of three tags used (points that were not visited are shown in grey). To test the effects of larger and irregular spacing between receivers on localisation error using multilateration and fingerprint localisation, two subsets of receivers (circles and triangles) were used.

To compare the multilateration and fingerprinting methods, 625 test points were generated along a 25 m x 25 m grid. For this test, we used solar/battery powered radio tags (CTT HybridTag, Cellular Tracking Technologies, Rio Grande, New Jersey, USA) that were programmed to signal every 5 seconds. Three tags were used at each test point. All tags were previously unused prior to testing. Tags were mounted atop a 2 m PVC pole (n = 16 poles, 48 tags). Two tags were mounted horizontally in opposite directions and one tag was attached vertically to the pole with the antenna pointed down. At each test point, a tag pole was held perpendicular to the ground for two to three minutes, and the pole was horizontally rotated 90 degrees approximately halfway through the test interval. While at the test point, the pole number, test point number, the start time (as the current minute, excluding seconds), and the end time (as the current minute, excluding seconds) were recorded. Each set of three tags was used at a row of 24 consecutive test points in order to distribute the tags throughout the study area in order to account for possible variation in tag signal strength (Figure 1). Additionally, at each point, the vegetation density was qualitatively scored as: 1) open field with no obstructions, 2) open with few shrubs nearby, 3) forested with no understory, 4) forested with little understory, 5) forested with thick understory.

We conducted two test sessions, the first on 23 March 2022 and the second on 23 June 2022, thus before and after foliage had fully emerged. The receiver grid was dismantled after the first test and then reinstalled and as a result the location of each receiver changed slightly (< 10 m) between the two tests. The same receiver, however, was used at the same location during the two tests to control for possible variation in sensitivity between devices.

### ii. Data processing

Following the tests, all data was retrieved from the base station as well as downloaded directly from the receivers to ensure that all detections were recovered. Duplicate detections were then removed and the data were filtered to retain detections from tags during the test intervals associated with the corresponding test point. As the test interval was only recorded as the current minute, the start and end times were rounded up and down, respectively, to the nearest minute and only detections within this reduced interval were retained to ensure that detections were recorded while the pole was stationary at the test point. All RSS values from each detection within each test interval were then averaged to give a single value for each receiver per tag. The full processed dataset included the following columns: test point, receiver ID, tag ID, date time, and RSS value.

### iii. Multilateration localisation

RSS-based multilateration was first applied to predict tag locations. Multilateration uses an empirically derived relationship between RSS and distance to estimate the distance, rather than the angle, between a signalling tag and the receiver. The relationship between RSS and distance for our tags and receivers was estimated by holding two tags mounted to a pole (as above) at set distances (1, 2, 5, 10, 15, 20, 25, 50, 75, 100, and 150 m) from two receivers for three minutes at each calibration distance. Non-linear least squares were then used to fit an exponential decay model:

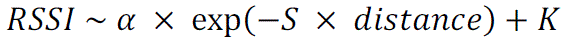

where `α` is the intercept, *S* is the decay factor, and K is the horizontal asymptote, as in (Paxton et al., 2022). Using the detections from our test tags and receivers, we estimated the model parameters as: `α` = 44, *S* = 0.01545, and K = –115, and these parameters were used in the multilateration localisation estimates (described below).

To optimize the multilateration estimate, we applied the RSS filtering approach of Paxton et al. (2022). This method first identifies the receiver with the strongest RSS, which is assumed to be the closest receiver to the signal, and then selects any other receivers that are within a set distance threshold. The distance threshold is arbitrary but should relate to the receiver grid spacing, with smaller thresholds minimizing the multilateration localisation error (Paxton et al., 2022). We applied a stringent filter of 1.5x the average receiver spacing (100 m). As such, receivers within 150 m of the receiver with the strongest average signal were retained and the RSS values from all detections were averaged for each receiver. We then used a log-distance path loss model (Seybold, 2005), implemented in the functions ‘ipfProximity’ and ‘ipfEstimate’ in the package *ipft* in R in order to estimate the position of the tag. The default values in the ‘ipfProximity’ function were then set to values found in the exponential decay model (above) of the RSS-distance relationship we measured with our tags and receivers.

Specifically, *alpha*, the path loss coefficient, was set to 4.17 based on the decay factor S of 0.01545 and the formula: alpha = –ln(S); *wapPow1*, the average RSS of a tag at one meter from a receiver, was set to –44, and *rssirange*, the observed range of RSS values was from –41 to –115. Lastly, the function ‘ipfEstimate’ was used to estimate the location of each of the three tags at each test point and the distance (in meters) between the estimated location of a tag and the true location of the test point was taken as the error.

### iv. RSS fingerprint localisation

RSS-based fingerprint localisation is a two-step process. In the first step, a radio map is created by measuring the RSS fingerprints at each test point (n = 625). The RSS fingerprints at each test point in consisted of the average RSS values from each tag registered by each receiver. Together these fingerprints comprise the radio map. In the second step, a classification algorithm is used to predict the location (coordinates) of RSS fingerprints by matching the fingerprint to the radio map. For this step, we used K-nearest neighbours (KNN) classification (Cover & Hart, 1967), an algorithm that is commonly used for indoor localisation applications (Obeidat et al., 2021). This method estimates the location of the test fingerprint by averaging the known locations of the K-nearest fingerprints in the radio map. KNN (k = 3) classification was performed using the ‘ipfKnn’ and ‘ipfEstimate’ functions from the *ipft* package in R (Sansano et al., 2019). As we had no prior expectation of the ideal choice of k and exploratory analysis suggested that different values did not impact the localisation accuracy, we used the default value (3) from the ‘ipfKnn’ function and the Euclidean distance between fingerprints was calculated to determine similarity. For each test point, the model was first trained on the remaining points (i.e. the radio map consisted of all points except the test point) and the location of each of the three tags at the test point was predicted. The localisation error was then calculated as the distance between the true location of the test point and the estimated locations of the three tags to compare this method against multilateration.

### v. Localisation error comparisons

We evaluated the performance of the multilateration and fingerprinting methods by comparing the median localisation error of each method. We also evaluated how features of the detection data that are commonly found in real-world tracking datasets affected localisation error. Specifically, we assessed the effect of not being detected by the nearest receiver to the test point, which could be caused by physical interference or by equipment failure. We also examined how features of the receiver network configuration affected localisation accuracy by simulating reducing the density and regularity of the receiver grid, both of which have been shown to negatively affect localisation accuracy for multilateration estimates (Paxton et al., 2022). To do so, we tested how using only half of the receivers in the grid would affect localisation accuracy (i.e., two subsets of receivers were used, Figure 1). Using only alternating receivers from the grid increased both the distance between the receivers and increased the variation in the inter-receiver distance (i.e., decreased the regularity of the receiver grid). Additionally, we evaluated how the test location within the study area affected the localisation accuracy for each method by examining the effects of distance to the nearest receiver, distance from the centre of the receiver grid, and vegetation density at the test point.

For fingerprinting, we also examined some situations that uniquely apply to this method. First, we evaluated the effect of removing different amounts of receivers randomly, 0.05 to 0.75 of all receivers, from the testing dataset. This process was repeated 50 times for each number of receivers removed. All receivers in the training data were, however, retained. As such, we effectively assessed how radio fingerprinting would be impacted by random equipment failure. We also evaluated the effect of varying amounts of fingerprinting time (5 to 60 seconds) at each test point on the error. Finally, to assess the robustness of the radio map to seasonal changes in vegetation, we used the radio fingerprints generated by the test points from the first round to localize tags at each test point in the second round.

In addition to KNN, other machine learning classification algorithms can be used to predict fingerprint locations and may yield better performance depending on features of the dataset, e.g., the number of receivers. As such, to examine whether other machine learning approaches showed marked improvement over KNN, we tested three other commonly used algorithms: random forests, XGBoost, and support vector machines (SVM), as well as a range of values of K (3-10) in KNN (Appendix A).

All data processing and analyses were conducted in the R environment (R Core Team, 2022). Code and data to recreate the analyses are available on GitHub at: https://anonymous.4open.science/r/radio_fingerprinting-DE51/README.md.

## Results

To compare the multilateration and fingerprinting methods, 532 test points were conducted (not all planned test points were accessible due to fences, water, and roads). Using multilateration localisation, it was possible to estimate tag positions at 461 of these test points since this method requires that at least 3 receivers detect the tag during the test interval, which did not always occur. As three tags were used at each location, this resulted in 1,376 and 1,596 test localisations for multilateration and fingerprinting, respectively. From these localisations, the overall median error was for multilateration 40 m and 30 m for fingerprinting (Figure 2). For comparison, we also applied a simple, proximity-based localisation method where the tag location is assigned to the receiver that detects the tag most strongly. Using this method, the median localisation error was 55 m.

**Figure 2.**
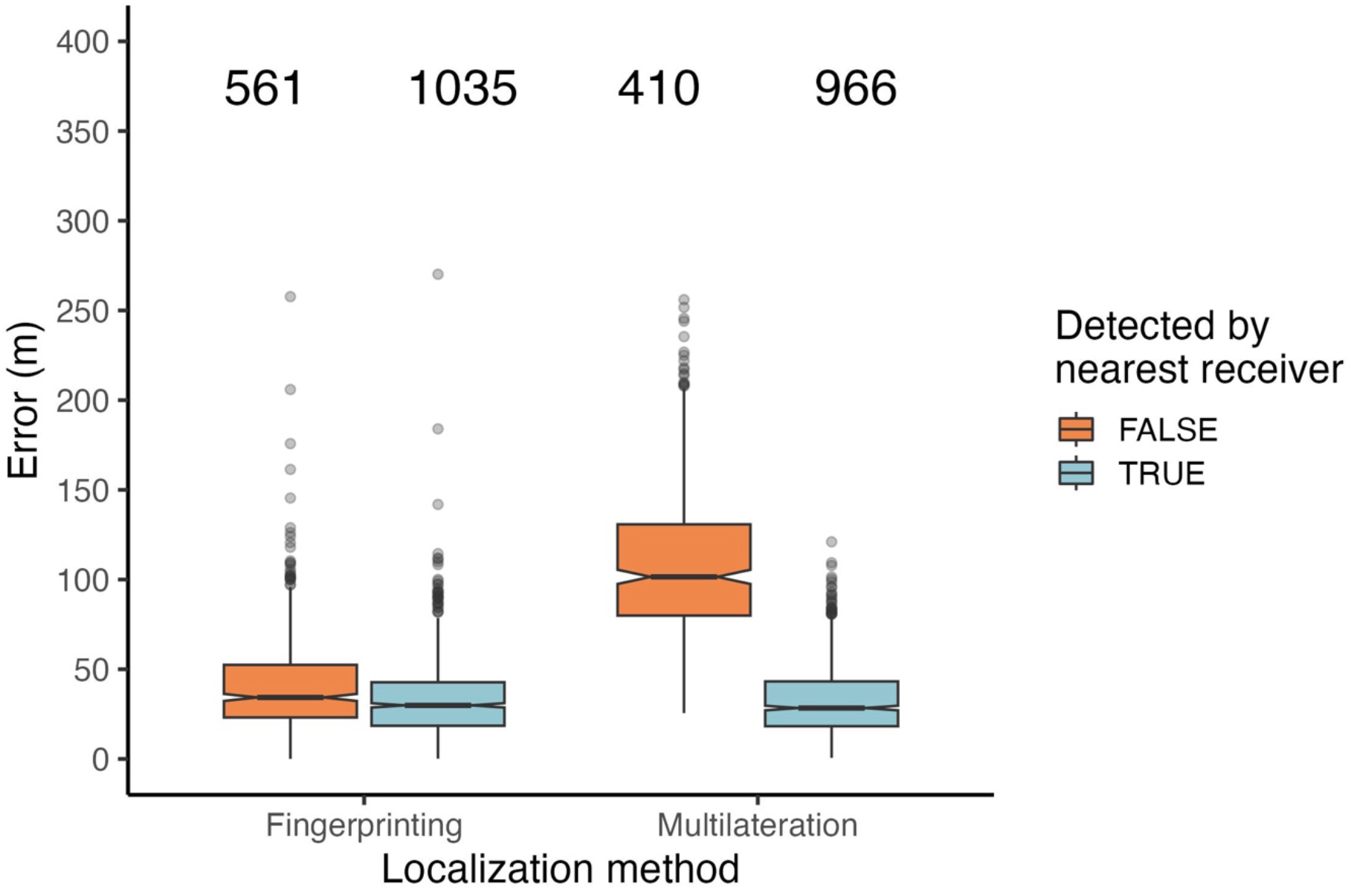
Notched boxplots of the localisation error (the difference between the true and estimated positions) for the multilateration and fingerprint localisation methods if the nearest receiver to the test point location did or did not detect the tag during the test interval. Median values are indicated by the centre of the notch and the notches correspond to the 95% confidence interval. Lower and upper hinges reflect the 25^th^ and 75^th^ percentiles and whiskers indicate values 1.5 times the inter-quartile range above and below the hinges. Outliers are shown individually as points. The number of localisations in each category are shown above each boxplot. The median localisation error was not significantly different between the two methods if the nearest receiver detected a tag at the test point (Wilcoxon signed-rank test, Z = –0.51, p = 0.57). When the nearest receiver did not detect the tag, however, the median error was significantly lower for fingerprinting compared to multilateration (Wilcoxon signed-rank test, Z = –23.29, p < 0.001).

Features of the test location affected the accuracy of each localisation method. In particular, whether the nearest receiver detected a tag at a test point strongly affected the median error for multilateration localisation. For the subset of multilateration localisations where the nearest receiver detected the tag (n = 966 of 1,376), the median error was significantly lower at 29 m as opposed to 101 m. In contrast, the fingerprinting median error was not affected by whether the nearest receiver detected the tag (Figure 2). Additionally, the test location was strongly related to median error for multilateration, but not fingerprinting, as the error between the estimated and true test location increased as the distance to the nearest receiver increased (Figure 3). Additionally, for multilateration compared to fingerprinting, the median error was lower when the test location was close to a receiver and higher when the test location was >40 m from the receiver. Median error for neither method, however, was significantly affected by the distance from the test location to the centre of the receiver grid (Figure 3). The median error for multilateration localisation estimates increased with higher vegetation density, particularly for the two most dense categories. For fingerprint localisation estimates, however, the median error was less affected vegetation density (Figure 4). When simulating removing alternating receivers from the entire grid and using the detections from only these two subsets (Figure 1, circles and triangles), the median error increased from 40 m to 60 m and from 30 m to 42 m for the multilateration and fingerprinting methods, respectively (Figure 5).

**Figure 3.**
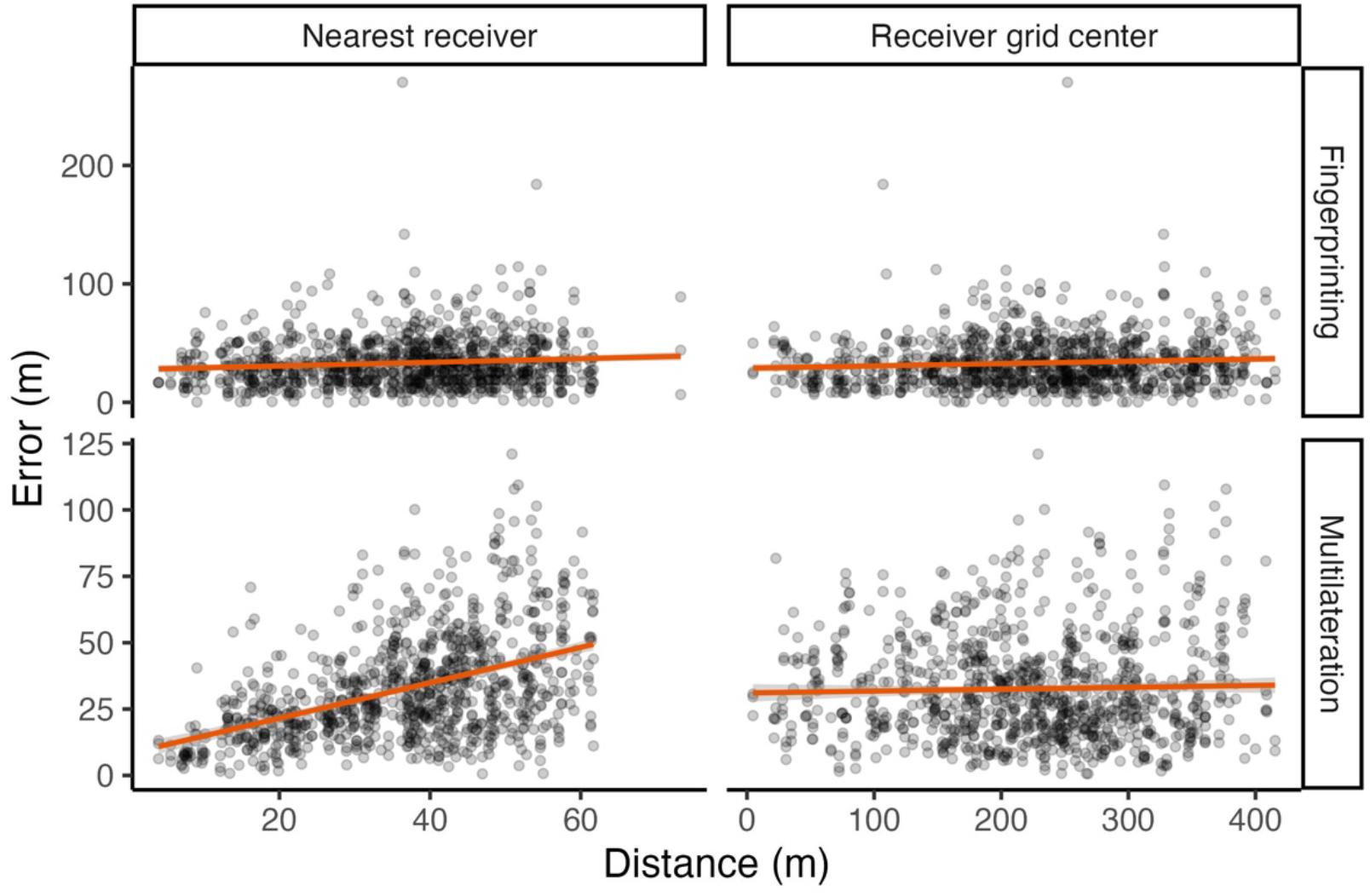
Error for fingerprinting (top row) and multilateration (bottom row) as a function of the distance to the nearest receiver (left panels) and to the grid centre (right panels) for localisations where the nearest receiver detected the tag. Orange line and shading represent the linear model regression slope and standard error.

**Figure 4.**
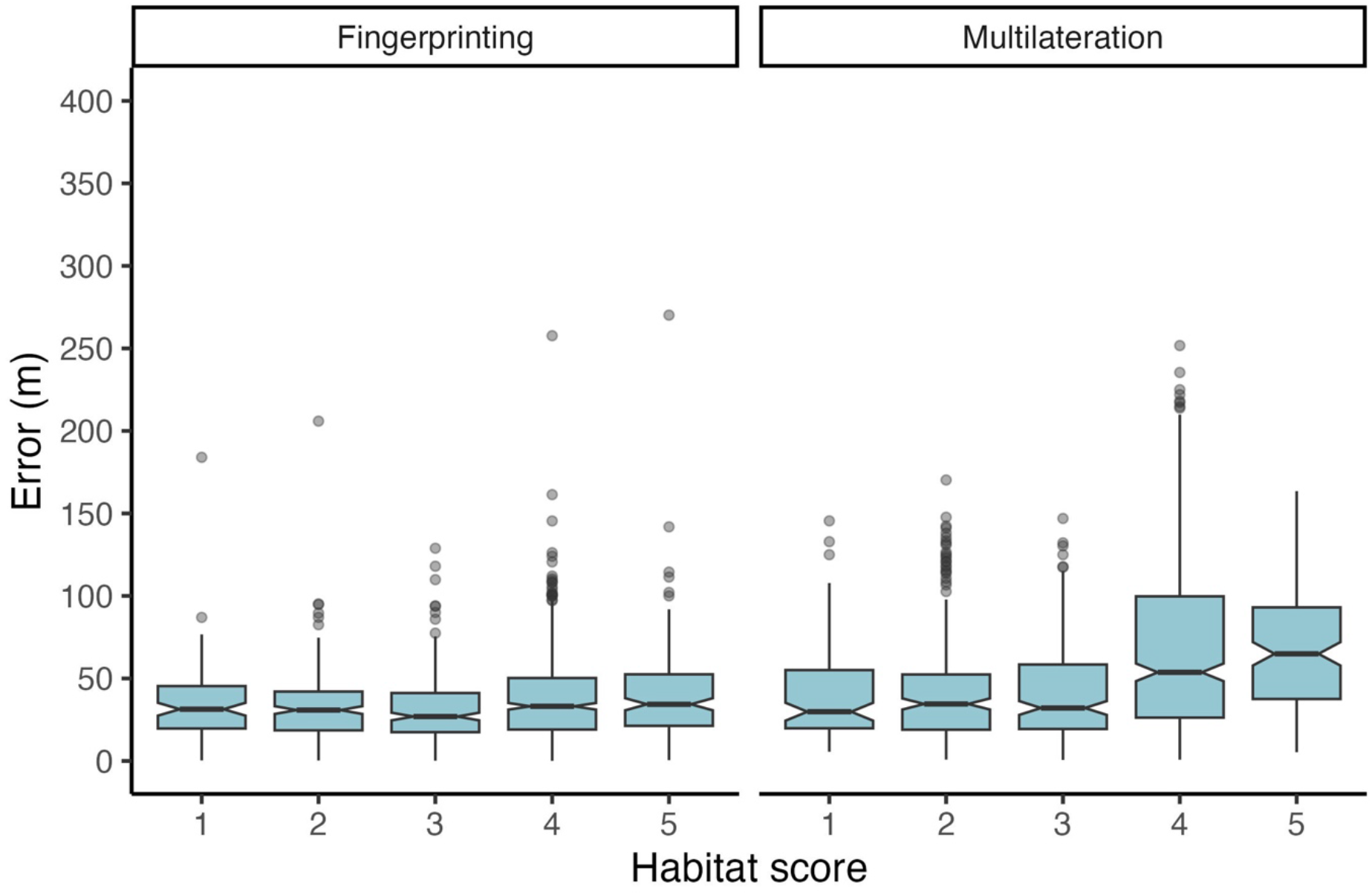
Notched boxplot of the error for each localisation method relative to the qualitative habitat score reflecting vegetation density at each test point. Median values are indicated by the centre of the notch and the notches correspond to the 95% confidence interval. Lower and upper hinges reflect the 25^th^ and 75^th^ percentiles and whiskers indicate values 1.5 times the inter-quartile range above and below the hinges. Outliers are shown individually as points. Vegetation scores correspond to following descriptions: 1) open field with no obstructions, 2) open with few shrubs nearby, 3) forested with no understory, 4) forested with little understory, 5) forested with thick understory.

**Figure 5.**
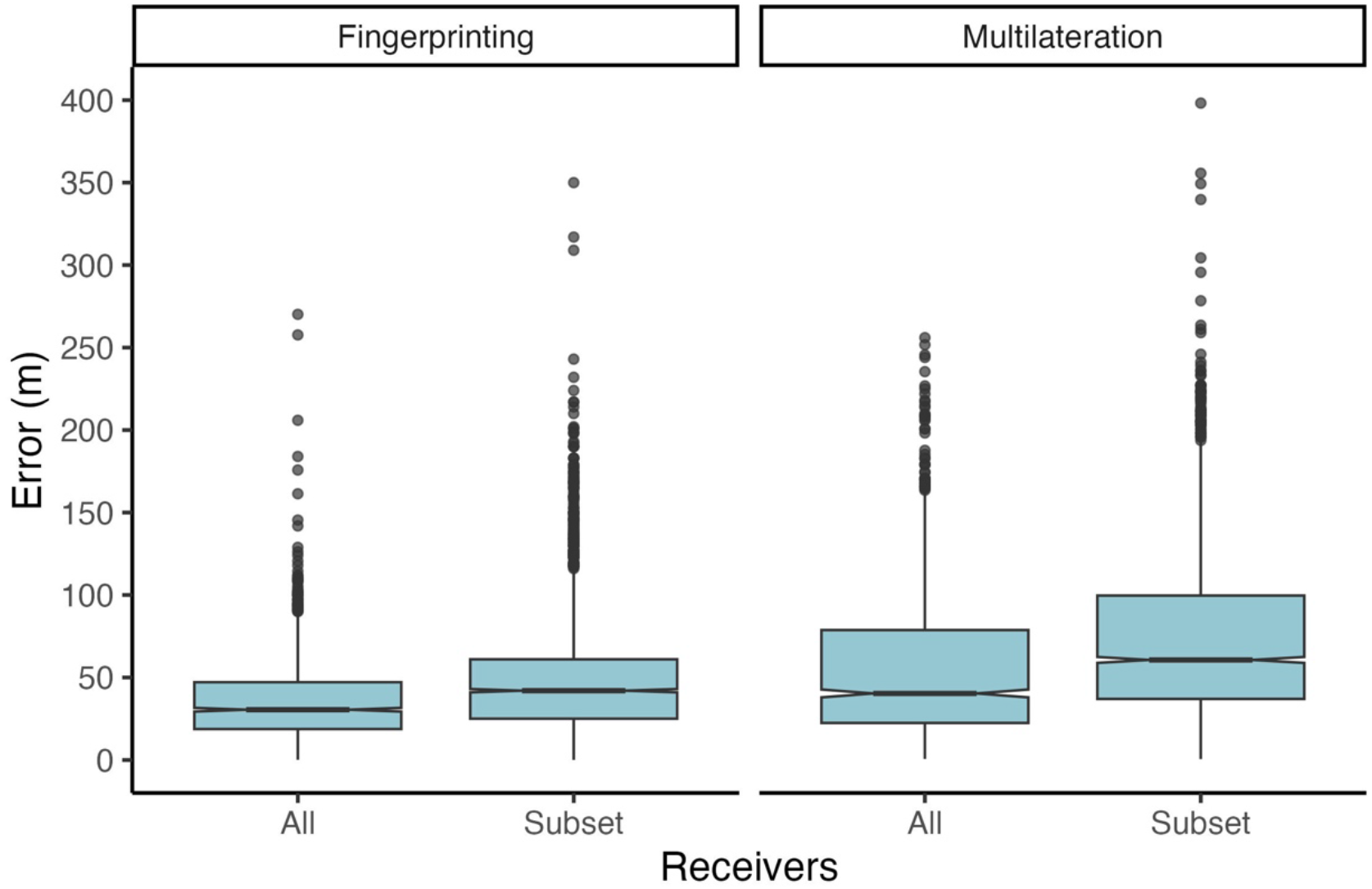
Notched boxplot of the localisation error when using all receivers or a subset of half of the receivers (as shown in Figure 1). Median values are indicated by the centre of the notch and the notches correspond to the 95% confidence interval. Lower and upper hinges reflect the 25^th^ and 75^th^ percentiles and whiskers indicate values 1.5 times the inter-quartile range above and below the hinges. Outliers are shown individually as points.

Reducing a random proportion of the possible receivers in the fingerprinting data did reduce localisation accuracy. The median error with reduced fingerprint receivers, however, only became comparable to the median error for multilateration localisations where the nearest receiver was not detected (i.e., 97 m) if only ∼0.65 of receivers were used (Figure 6). For fingerprint localisations, reducing the time spent at each test point significantly decreased the detections registered per tag with the effect being more pronounced in more densely vegetated habitats (Figure 7a). Relatedly, reducing the time spent at each test point increased the median error, though only up to 20 seconds when the median error was minimized and additional time at the test point only marginally reduced the median error (Figure 7b). The same general relationship between time spent at the test point and median error was observed across habitat types. The median error from each of the machine learning algorithms, except XGBoost, was lower than that of multilateration (Figure A1).

**Figure 6.**
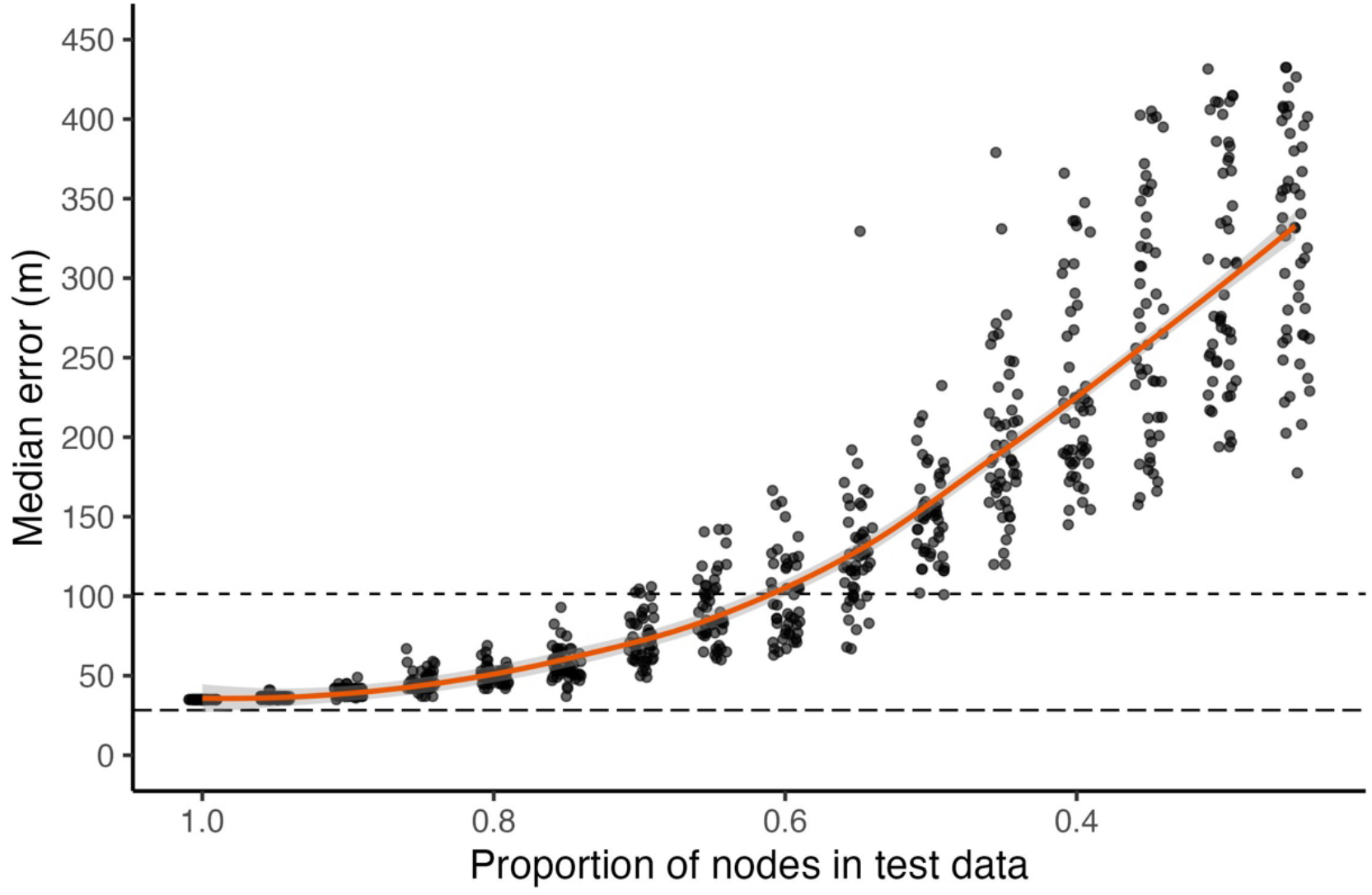
Relationship between the proportion of receivers retained in the radio fingerprint testing data used for the fingerprint localisation estimates. Random receivers, ranging from 0 to 0.75, of all receivers in the training dataset were removed from the fingerprint testing dataset and the test fingerprint position was predicted. The locally weighted linear regression (LOESS) fit is shown for reference. The long-dash line shows the overall median error for multilateration localisation and the short-dash line shows the median error when not detected by the nearest receiver for multilateration localisation.

**Figure 7.**
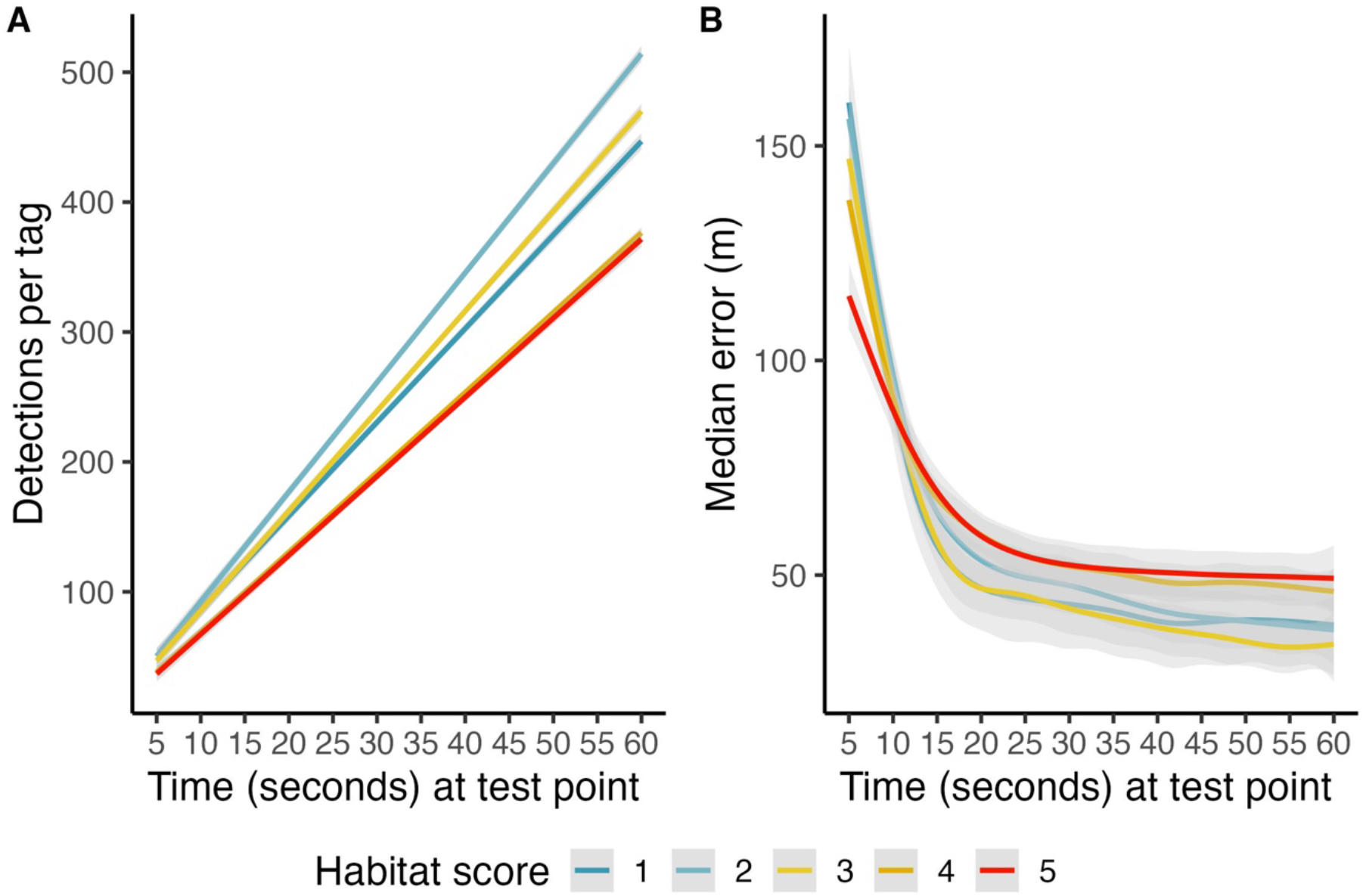
Relationships between the time (seconds) at each test point for subsets of calibration data for points in each habitat group and A) the total number of detections per tag and B) the median error (m) from all fingerprint localisations. Habitat scores correspond to a qualitative evaluation of the amount of vegetation at the test point: 1) open field with no obstructions, 2) open with few shrubs nearby, 3) forested with no understory, 4) forested with little understory, 5) forested with thick understory vegetation.

Lastly, predicting tag locations during the second test using the radio map constructed from detections during the first test increased the median localisation error from 30 m to 50. Applying the radio map from the second test to predict tag locations from the second test, however, resulted in a similar, but slightly lower median error of 25 m compared to the first test. The median localisation accuracy for multilateration during the second test was 38 m (compared to 40 m during the first test).

## Discussion

We have compared the accuracy of two general methods, multilateration and fingerprint localisation, for estimating radio tag positions in automated radio tracking systems. Such systems are being increasingly used to track animals that are range resident within areas of moderate spatial extent (generally <5 km^2^) owing to the relative simplicity of installation and the ability to affordably track many individuals simultaneously with high temporal resolution over extended periods. We found that while the overall median error was similar between the two methods, location estimates using the fingerprinting method were more robust to multiple scenarios that are likely to occur in real-world tracking studies. While previous studies have used machine learning approaches to localize radio tags, these studies differed significantly, predicting locations along a single dimension (Harbicht et al., 2017), utilizing only a few receivers to produce relatively course location estimates (Scardamaglia et al., 2022), or within a small area (0.04 hectares, Wallace et al., 2022). Here, we have demonstrated fingerprint localisation as an effective method to obtain high-frequency, fine-scale localisation estimates of track animals using automated radio telemetry systems.

Aside from evaluating radio fingerprinting as localisation method in an outdoor, natural environment, we assessed how various conditions relating to the test location and to the receiver grid configuration affected the localisation error of the two methods. Overall, the median error was similar between multilateration (28 m, if detected by the nearest receiver) and fingerprinting (30 m). For multilateration, the error we observed in our field tests is similar to the lower values reported by Paxton et al. (2022) which simulated a receiver grid with 100 m spacing and evaluated the effect of different filtering methods for multilateration localization. This suggests that the error we observed using multilateration in this field study is close to the minimum feasible given the receiver configuration we tested. Multilateration localisation error, however, increased nearly 3.5-fold in cases where the nearest receiver to the test point did not detect a tag. This could be caused by an inactive receiver or by interference from the environment, e.g., vegetation or other solid structures, all of which could prevent the tag signal from being received. Missed detections are likely to occur in field tracking studies and would be impossible to detect when estimating the unknown location of a tagged animal. As such, our results reveal an important limitation of multilateration approaches relative to fingerprint localisation methods. Localisation error was also not consistent throughout the study area as it increased with distance from the nearest receiver and in more densely vegetated areas. These findings highlight that localisation accuracy for range-based methods is dependent on signal strength and that factors which limit signal strength, such as pockets of dense vegetation, increases the localisation error when using multilateration (Whitehouse et al., 2007).

Receiver configuration and placement is an important consideration when designing an automated radio telemetry system since for a given number of receivers, decreasing the receiver density increases the spatial extent that is covered, but potentially at the cost of reduced localisation accuracy. In our test, we made use of a relatively dense receiver network in which receivers were spaced every 100 m, which was the lowest receiver spacing tested by Paxton et al. (2022) where the effects of receiver distance and spacing regularity on localisation accuracy. We similarly explored how accuracy using multilateration and fingerprint localisation was affected by using two subsets of the receiver network (Figure 1) which resulted in an irregular grid where each three nearest neighbours for each receiver were 100 m to 200 m away. While median error increased for both localisation methods, the change was more pronounced for multilateration, increasing by 50% (Figure 4). These results indicate that using fingerprint localisation allows for greater flexibility in receiver grid construction as regular spacing is both less critical and receivers can be spaced farther apart to increase the extent of the network without substantially affecting the localisation accuracy. Given that in complex habitats where certain areas are inaccessible due to terrain, positioning receivers in a regular grid can be difficult, and so greater flexibility in receiver placement will be a practical benefit of using fingerprint localisation.

While our results indicate that fingerprint localisation offers several benefits over multilateration, there are additional aspects to consider when applying fingerprinting methods. One consideration is the construction of the radio map. For indoor positioning applications, extensive research has focused on radio map construction methods and their limitations (Han et al., 2014; Jung, Lee, et al., 2017; Jung, Moon, et al., 2017). In this study, we applied a point-by-point manual calibration procedure, which is the most time intensive radio mapping method, but typically produces the highest localisation accuracy (Jung, Moon, et al., 2017). In practical terms, the time spent at each calibration point will strongly determine how long the radio mapping process takes. In this study, we observed, however, that reducing the amount of time spent at each calibration point to 15 seconds from 60 seconds did not substantially affect the accuracy using the fingerprint localisation method (Figure 7B). Another radio map construction method that can yield similar performance to point-by-point manual calibration is the walking survey (Jung, Moon, et al., 2017). This method only involves specifying routes for the radio mapping such that the receiver network is sufficiently covered to achieve the desired localisation accuracy which is determined by the density of the survey routes. Another consideration to applying a fingerprint localisation approach is that if the environment is significantly altered, through, for example, seasonal changes in vegetation, or if the receiver network changes, then the radio map may no longer be representative (He & Chan, 2016). Our initial test occurred during boreal winter prior to spring green-up while our second test occurred during boreal summer when vegetation density was significantly higher. Additionally, receiver placement was not identical in both tests. These factors would be expected to impact the local radio fingerprint and in the second test, the localisation error increased to 50 m when predicting tag locations from the fingerprints collected during the second test using the initial radio map (though the median error dropped slightly to 25 m when using the radio map constructed during the second test, which was likely due to larger differences in radio fingerprints owing to increased local interference from vegetation). Adapting to changes in the radio map is an important consideration for effective fingerprint localisation. While there are a variety of solutions to the problem from the field of indoor positioning, which generally involve monitoring signal changes through additional infrastructure, such as using stationary beacon tags and updating the radio map if localisation accuracy degrades (He & Chan, 2016), this may not always be feasible. Consequently, while fingerprint localisation has several advantages over multilateration, this method may not be suitable for every application and so optimized multilateration approaches, such as those developed in Paxton et al. (2022), are also highly valuable.

Aside from the radio map construction method used, another consideration for fingerprint localisation is the choice of a suitable machine learning algorithm for the classification of fingerprint locations (reviewed in Roy & Chowdhury, 2021). Our objective in this study was to compare multilateration and fingerprint methods for radio tracking in an outdoor, natural environment, rather than an extensive evaluation of the performance of different classification models. We did, however, consider the performance of four other commonly used machine learning algorithms (Appendix A) and all but one (XGBoost) produced similar median errors, which were lower than that obtained by multilateration. We focused on KNN as variants of this method are widely used since they are easy to implement as only two parameters must be specified, the number of neighbours (k) and the distance function. While we found that this method performed well, there is extensive ongoing research into developing learning techniques to optimize localisation estimates with radio fingerprinting. Where large fingerprint datasets can be collected, methods such as Deep Learning, may be used to minimize localisation error.

Automated radio telemetry systems are continually developing and are already being employed to study diverse topics ranging from social networks in birds (Shizuka et al., 2022; Snijders et al., 2014) to reintroduction techniques for conservation programs (Smetzer et al., 2021) to habitat selection in flying insects (Fisher & Bradbury, 2022). Given the advantages of this tracking method, such as applicability, duration, and temporal resolution, it is expected to remain an important tool in wildlife tracking studies (Nathan et al., 2022). By applying suitable analytical techniques, such as the radio fingerprinting method we have presented in this study, researchers will be able to further realize the full potential of these tracking systems.

## Acknowledgements

We are grateful to Geldersch Landschap & Kasteleen for permission to conduct fieldwork in Westerheide Wildlife Sanctuary. Also, we thank Andrea Tribulato, Andries van Vuuren, Bryan Fitzmaurice, Caroline Heylen, Catherine Nistha, Clara Vinyeta Cortada, Elke Molenaar, Elza van Delft, Frank Rooijakkers, Gijs van der Gun, Gwyn Hietbrink, Job Hooft, Jori Nordenboos, Judith de Veer, Lynn van Boheemen, Maria Fernandes Rosas Corona, Novi Glissenaar, Paja Biglova, Pien van der Poll, Puck Heijmink, Sarinda Westerhout, Shareena Dwarka, Subin Santosh, Thomas van der Heijde, and Ziyu Li for help collecting data during the field tests. C.T. and H.Y. were funded by Next Level Animal Sciences innovation initiative of Wageningen University & Research. E.S.S was funded by the Spanish Ministry of Science and Innovation through Project INDRI under Grant PID2021-122642OB-C44. Additional funding for this project was provided to C.T. by the Wageningen Data Competence Centre Data Science and Artificial Intelligence 2022 Fellowship program.

## Author Contributions

C.T. conceived the ideas and designed the methodology; C.T. and R.F. collected the data; C.T., H.Y., and E.S.S. analysed the data; C.T. and M.N. led the writing of the manuscript. All authors contributed critically to the drafts and gave final approval for publication.

## Data Availability

R code and data to reproduce analyses from this study and to facilitate radio fingerprint localisation are available on GitHub at: https://anonymous.4open.science/r/radio_fingerprinting-DE51/README.md. The radio fingerprinting dataset is available at https://zenodo.org/records/10026887.

