## Appendix A for "Fingerprint localisation for fine-scale wildlife tracking using automated radio telemetry"

### Appendix S1. Comparison of machine learning algorithms for fingerprint localisation

Fingerprint localisation entails matching new radio fingerprints to points in the radio map based on similarity in order to estimate the location of new radio fingerprints. To identify the most similar points, machine learning algorithms are typically used. While nearly any machine learning algorithm can be used, various factors such as the number of radio receivers, the amount of environmental noise, and the complexity of the training dataset, will affect the suitability of different approaches. KNN is considered to be one of the most robust machine learning approaches for fingerprint location classification, but other approaches, such as random forests, Support-Vector Machine (SVM), and XGBoost are suggested to potentially perform better in noisy, irregular environments (Bozkurt et al., 2015; Nessa et al., 2020). We compared the localisation accuracy of each these methods as well as KNN with varying numbers of neighbours (3 to 10).

To compare each of the machine learning methods, the focal test point was withheld from the training data and the location of each of the three tags at the test point was predicted. The implementation of KNN was described in the main text. For random forests, SVM and XGBoost, regressions on longitudinal and latitudinal directions were preformed separately. For example, in the case of random forests, two individual models (i.e., on longitude and latitude data) were trained and tested on each train-test combination. Then the localization error was calculated as the distance between the true location and the predicted location of the test tag.

Random forests were fit using the *ranger* package in R based on a 70% split of the data (Wright & Ziegler, 2017). Hyperparameters were tuned using a grid search for the number of randomly selected predictors at each split (parameter: mtry, values: 2 to 10), the splitting rule (splitrule, values: variance, extratrees, maxstat, beta), and the minimum node size (min.node.size, values: 5 to 10) using the function ‘train’ from the R package *caret* (Kuhn, 2008). The parameter values selected for model fitting were: 9 (mtry), variance (splitrule), and 5 (min.node.size). SVM models were fit using the *e1071* package (version 1.7-9) in R (Dimitriadou et al., 2006). The default settings of *svm* function were used where a radial kernel was applied. XGBoost models were fit using *xgboost* package (version 1.5.2.1) in R. The “nrounds” was set to 100 and other hyperparameters were kept as defaults using the ‘xgboost’ function (Chen & He, 2023).

Overall, each of the machine learning algorithms performed similarly (median error, 28 to 36 m) with the exception of XGBoost, which had a higher median error (43 m). For KNN, the choice of K did not strongly affect the median error as there was only a 2 m difference between the lowest median error (K = 10) and the highest median error (K = 4).

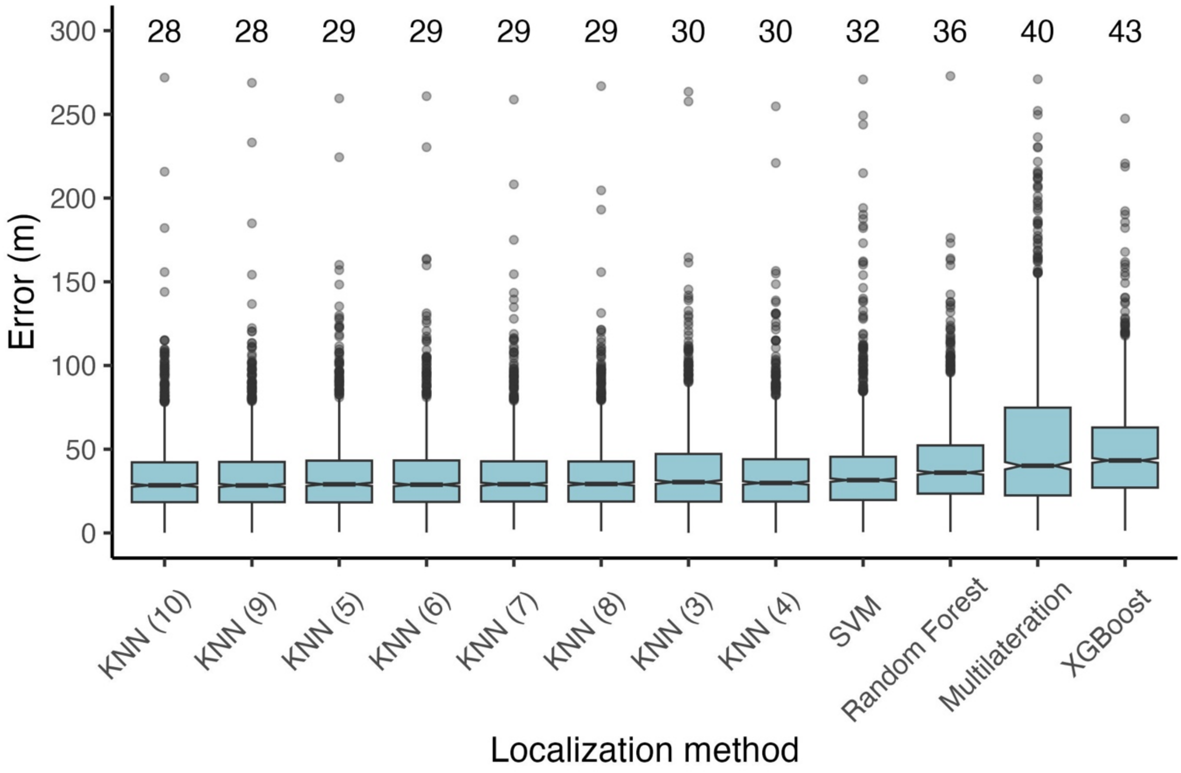


Figure S1. Notched boxplot of the error (m) for each localisation method examined. The methods are arranged by median error in ascending order (median error shown above each boxplot). Median values are indicated by the centre of the notch and the notches correspond to the 95% confidence interval. Lower and upper hinges reflect the 25$^{th}$ and 75$^{th}$ percentiles and whiskers indicate values 1.5 times the inter-quartile range above and below the hinges. Outliers are shown individually as points. Non-overlapping notches between two groups indicate that the medians in those groups are significantly different.
